# A landscape description of the dynamics of Turing patterns

**DOI:** 10.1101/2025.06.13.659478

**Authors:** Shubham Shinde, Archishman Raju

## Abstract

Turing patterns are a well-studied model of reaction-diffusion equations for developmental patterning. Their applicability has often been limited by the difficulty in identifying candidate molecules that satisfy the requisite criteria for patterning. Here, we build on recent work on geometric models to describe Turing patterning as a potential flow. We show how the universal dynamics of Turing patterning is described by a landscape, largely independent of the underlying reaction-diffusion equations. We apply our framework to three-component systems and demonstrate that we can accurately capture the dynamics of any given component. We extend our framework to larger networks and to models of Turing patterns coupled with external morphogens that provide positional information. We provide a quantitative description of the dynamics of chosen markers and apply it to the dynamics of SOX9 expression during digit patterning.

## I. INTRODUCTION

Turing patterns are one of the oldest models for spatial pattern formation in the context of developmental biology [1]. The most well-known mechanism of Turing patterns is the activator-inhibitor system [2]. Despite some success in identifying candidate activator inhibitor pairs [3–5], it has generally been difficult in developmental systems to identify candidate molecules whose regulatory connections and diffusivities satisfy the requisite criteria for Turing patterns [6–8].

The proposal that a three-component Bmp-Sox9-Wnt network could be responsible for the patterning of digits in mice has therefore excited considerable interest in the investigation of Turing patterns with three or more components, some of which may be non-diffusible [7, 9, 10]. Further, it has been suggested that the coupling of Turing patterns with other morphogen gradients can lead to more complex dynamics [11–13].

These models are all described by reaction-diffusion equations. The reaction terms are typically phenomenological, with different models making different choices. Parameter values like rate coefficients, thresholds, etc., are rarely known, and the complexity of signaling pathways is typically not modeled. It is now well known that models of this form suffer from over-parameterization and typical predictions are sensitive only to a few directions in parameter space [14]. There is therefore considerable interest in understanding the low dimensional behaviour of spatial patterns in developmental systems.

A substantial literature exists on approximating the dynamics of pattern-forming systems modeled using reaction-diffusion equations. It is common to apply a perturbation theory in the bifurcation parameter giving rise to the so-called amplitude equations [15–18], which provide an approximate quantitative description of the nonlinear dynamics. For Turing patterns on a finite do-main, it is possible to write ordinary differential equations for the evolution of a finite set of unstable modes. The method of normal forms uses near-identity coordinate changes to reduce this system of equations to the simplest form that qualitatively preserves the dynamics.

In practice, one may observe the concentration of a localized patch of one or more component molecules (e.g.morphogens) increase in time and saturate without full knowledge of all relevant regulatory components. Our increasing ability to measure such spatio-temporal dynamics of gene expression has led to an interest in quantitative models to describe this dynamics.

It was suggested in Ref. [19] that the quantitative dynamics of developmental systems can be described by a potential landscape with a Riemannian metric under quite general conditions. The potential landscapes are minimally parameterized and can be directly inferred from the data without knowledge of the underlying molecular constituents. These ideas have now been applied to a wide variety of developmental systems including *C. Elegans* vulval development [20], inner cell mass (ICM) specification in the mouse blastocyst [21], and differentiation of mouse embryonic stem cells (mESC) [22]. These geometric ideas have also been explored in multicellular contexts [19, 23–25].

The contribution of this manuscript is as follows. First, we use the mathematics of hypernormal form theory to simplify arbitrary reaction-diffusion equations into their simplest form. We show that these equations are well described by a landscape. We then give a specific prescription for how this landscape can be used to quantitatively describe observable expression. The landscape description not only captures the qualitative behaviour of patterns, but allows us to do a quantitative phenomenology of the dynamics. The landscape parameters can be directly inferred from data.

We demonstrate the utility of our method by comparing this landscape approximation to numerical solutions of various three and more component Turing networks. We also extend our framework to include Turing networks where a parameter is dependent on a spatially varying morphogen profile. Finally, we apply our framework to study the dynamics of SOX9 expression in the context of digit patterning.

## II. NORMAL FORM THEORY FOR REACTION-DIFFUSION EQUATIONS

Normal form theory provides a systematic analysis of the behaviour of ordinary differential equations. Normal forms eliminate terms at a certain polynomial order by near identity coordinate changes of the same polynomial order [26]. Normal forms are used to understand the qualitative structure of the flow. Hypernormal form theory goes a step further and uses lower-order polynomials to remove higher-order terms to reduce the equations to the simplest possible normal form [27]. For completeness, we show how these coordinate changes can be performed for reaction-diffusion systems on finite domains. We then show with an example how the resultant system can be written as a landscape with the help of a metric.

Turing patterns are modeled as reaction-diffusion equations. We assume that these equations can be written in the following general form

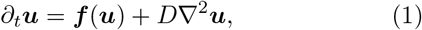

where ***u*** = [*u*_1_, *u*_2_,, … *u*_*N*_] are the concentrations of the components (e.g. gene expression), *D* is a diagonal matrix with diffusion constants (some of which could be zero), ***f*** (***u***) are reaction terms which are nonlinear in general and reflect regulatory and signaling interactions between the underlying components, and ∇^2^ is the Laplacian. We use periodic boundary conditions, unless mentioned otherwise. Henceforth, vectors are denoted using bold symbols.

Taylor expansion of the reaction terms in Eq. (1) about the homogeneous state ***u***_0_ gives

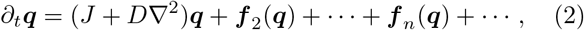

where ***q*** = ***u*** − ***u***_0_, *J* is the Jacobian matrix evaluated at ***u***_0_, ***f***_*n*_(***q***) are homogeneous polynomials of order *n*, taking the form 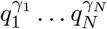, where the exponents satisfy *Σ*_*i*_*γ*_*i*_ = *n*. Since the Turing instability is diffusion-driven, the reaction terms must be stable around ***u***_0_. The linear stability analysis is obtained by substituting ***q*** = ***v****e*^*ikx*+*λt*^ in the linear part of Eq. (2), which leads to the eigenvalue equation *λ****v*** = (*J* − *Dk*^2^)***v***. There are *N* eigenvalues *λ*^(*i*)^(*k*) which are a function of the mode *k*. For small networks (*N* = 2, 3), at most one solution of the characteristic equation det |*J*− *Dk*^2^ − 𝕀*λ*_*k*_| = 0 exhibits an instability (Appendix A). More generally, even for larger networks close to a bifurcation, only one eigen-value has an instability, leading to a few unstable modes while other eigenvalues remain stable. We therefore focus on the dynamics along a single unstable eigenvalue *λ*^(1)^(*k*) and adopt the ansatz

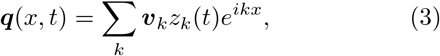

where *z*_*k*_ is the complex amplitude, and ***v***_*k*_ is the eigen-vector corresponding to the unstable eigenvalue. Since the solution ***q*** is real, the amplitudes satisfy 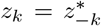. Substituting the above ansatz in Eq. (2) gives

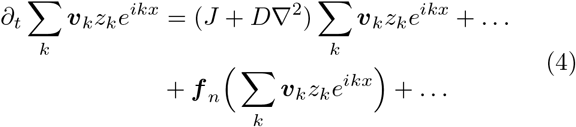

We project the above equation along the *k*-th mode using the operator (2*π*)^−1^ § d*xe*^*−ikx*^ followed by dot product with the left eigenvector corresponding to ***v***_*k*_. This gives the evolution equation for *z*_*k*_

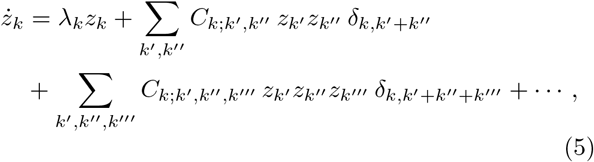

where, *δ*_*i*,*j*_ is the Kronecker delta, equal to 1 if *i* = *j* and 0 otherwise. Therefore, in the equation for *z*_*k*_, a term of order *n* remains only if the sum of its modes is equal to *k*. The coefficients 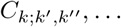 are real and depend on the nonlinear terms in the reaction kinetics. At linear order, we get the eigenvalue *λ*_*k*_ which is real. With two modes *k* = 1, 2 in the ansatz, including all allowed terms gives the equations for *z*_1_ and *z*_2_

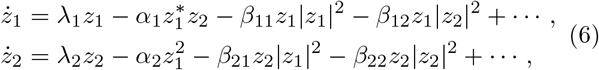

where we have denoted the coefficients at quadratic and cubic order by *α*_1_, *α*_2_ and *β*_11_, *β*_12_, … respectively for notational convenience. The equation above can alternatively be derived from symmetry arguments [28].

With *N* modes retained, Eq. (6) can be written in the general form

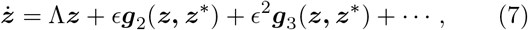

where ***g***_*n*_ are homogeneous polynomials of order *n*, Λ is a diagonal matrix with whose entries are eigenvalues *λ*_*k*_, and *ϵ* is a bookkeeping parameter to track the polynomial order. We treat ***z*** and ***z***^*^ as independent variables. Since the conjugate equations are identical, it suffices to consider only the equations for ***z***.

To simplify terms at polynomial order *p*, we perform a coordinate transformation 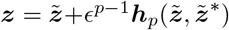. Here, 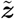 are the new coordinates, and ***h***_*p*_ is determined to remove as many terms as possible. Taking the time derivative of the coordinate transformation gives

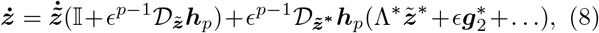

where 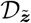 is the matrix of first-order partial derivatives with respect to variables 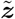, 𝕀 is the identity matrix, and we have substituted the conjugate of Eq. (7) for 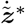. We use the expansion 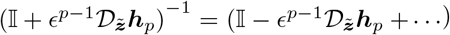 and substitute the time-derivative given by Eq. (8) into Eq. (7). Collecting terms at polynomial order *p* in the resulting expression gives

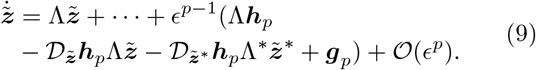

To remove terms at order *p*, we seek ***h***_*p*_ such that the *ϵ*^*p−*1^ order term vanishes. This condition can be written as [***h***_*p*_, Λ*z*] + ***g***_*p*_ = 0, where […] denotes the Lie bracket. This is a linear equation acting on a linear vector space with a natural basis formed by products of monomials and the standard basis vectors. Expanding both ***g***_*p*_ and ***h***_*p*_ in this basis, we associate each term *v****z***^***γ***^***z***\*^***δ***^***e***_*i*_ in ***g***_*p*_ with a corresponding term of the same monomial form *m****z***^***γ***^***z***\*^***δ***^***e***_*i*_ in ***h***_*p*_, where 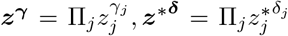, the exponents satisfy Σ_*j*_ (*γ*_*j*_ +*δ*_*j*_) = *p*, ***e***_*i*_ is the *i*-th standard basis vector and the constant *m* is real. Substituting these terms into [***h***_*p*_, Λ*z*] + ***g***_*p*_ = 0 gives the solution for

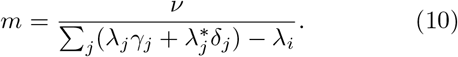

For 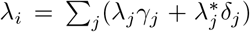 there is no solution for the above equation, and the corresponding term in ***g***_*p*_ cannot be eliminated. This defines the resonance condition.

After removing the non-resonant terms, we write Eq. (7) as 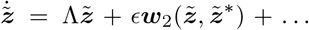, where ***w***_*p*_ are resonant terms of order *p*. We make a coordinate transformation 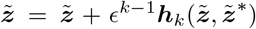 at order *k < p*. At order *p*, keeping only leading order terms in ***h***_*k*_, we get ***w***_*p*_ + [***h***_*k*_, ***w***_*p−k*+1_]. Thus, we can remove higher order resonant terms by solving ***w***_*p*_ + [***h***_*k*_, ***w***_*p−k*+1_] = 0. At a hyperbolic fixed point, all the nonlinear terms can be eliminated in the absence of resonances. However, we only allow coordinate changes that are non-singular at the bifurcation.

We demonstrate the above procedure for one unstable mode (*k* = 1) below. Keeping all terms that are resonant at *λ*_1_ = 0 in Eq. (6) gives the following equations

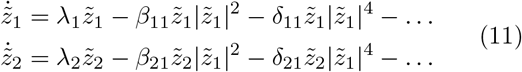

To remove the fifth-order terms in Eq. (11), we make a coordinate transformation at cubic order with 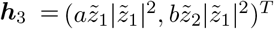 for real *a* and *b*. This gives [***h***_3_, ***w***_3_] = (0, −2*bβ*_11_)^*T*^, so the fifth-order term in the 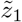 equation cannot be removed. Choosing *b* = −*δ*_21_*/*(2*β*_11_) removes 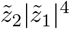 term from the 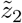 equation. All the higher-order terms can be removed, and we get the normal form

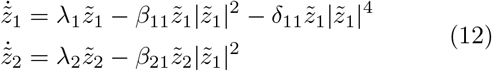

Although the normal form contains a fifth-order term, we show later that retaining terms only up to cubic order suffices for the applications that we consider. Mainly, we are interested in the dynamics of patterns that start near a homogeneous state and saturate. Therefore, the normal form equation along the unstable mode becomes

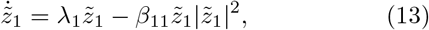

*λ*_1_ and *β*_11_ are parameters to be fit from the data. For a single unstable mode, the normal form can be trivially derived from a potential landscape. Though Eq. (13) commonly occurs as an approximation to the amplitude of an unstable mode, our interpretation of the equation is different. The equation is the simplest normal form that can be obtained by analytic coordinate changes (ignoring the 5th order term). We require coordinate changes to explicitly connect it to observable patterns as we do in the next section.

For the two unstable modes (*k* = 1, 2) case, in Eq. (6) we also keep terms only up to cubic order. Note that the coefficients of the cross-terms in Eq. (6) are not necessarily the same and depend on the dependence of the eigenvectors on the mode *k*. These equations are generically not in potential form. However, we can construct a genetric potential to quartic order including the allowed terms in Eq. (6). We show with an example in SI Section I that the dynamics of Eq. (6) and the dynamics generated by the gradient of the potential only agree with the inclusion of a suitable metric

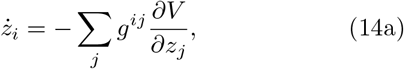

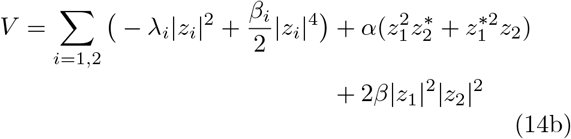

where, *g*^*ij*^ is the inverse metric tensor. In practice, we do not require the metric to fit the dynamics for reasons described in the SI and hence do not consider it further.

Intuitively, the landscape captures the geometry of the dynamics and is written in terms of the normal form coordinates. Consider the case of one unstable mode. In the original coordinates (Fig. 2A, black grid) the stable patterned state has a contribution from both modes. The coordinate transformation (Fig. 2A, blue grid) aligns the dynamics with one of the coordinates.

Given data on the concentration of a molecular component (Fig. 2B), we can project it onto the normal form coordinates and fit a universal form to the dynamics (Fig. 2C). Along with the dynamics of the normal form coordinate, we can also fit a time-independent coordinate change that captures the dependence of all other modes on the dynamics of normal form coordinates (Fig. 2C). As shown below, we can predict the general form of these coordinate changes using a perturbation theory.

## III. CORRECTIONS FROM COORDINATE CHANGE

To relate the dynamics in the normal form coordinates to the concentration of a molecule, we need a map from the normal form coordinates to the observable expression (Fig. 1). This map is obtained by coordinate changes.

**FIG. 1.**
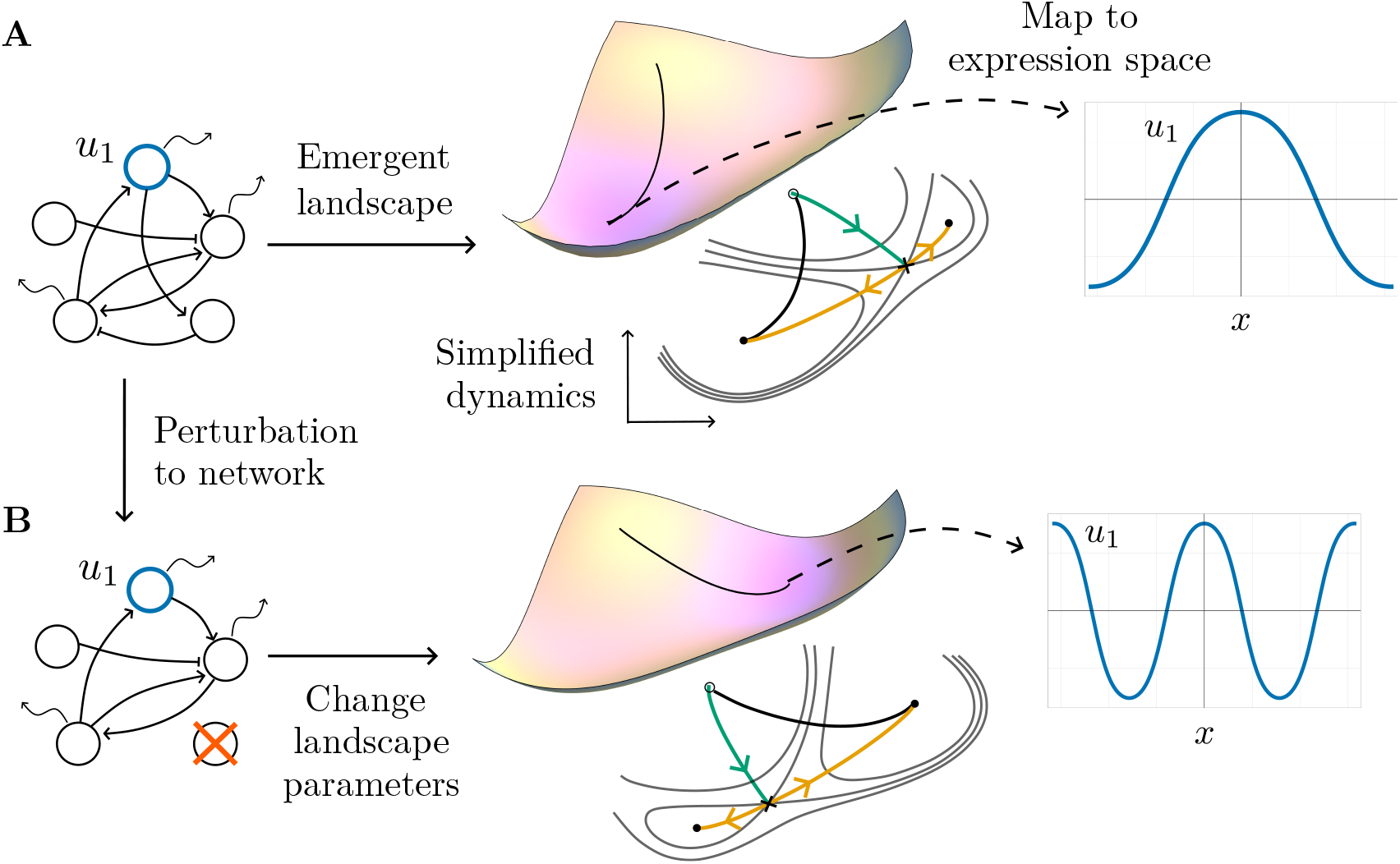
The dynamics of Turing patterns has an emergent landscape description. (A) Turing patterns are described by reaction-diffusion equations for a set of components that regulate each other’s activity (diffusible components are shown using wiggly arrows in the network). These components could be signaling ligands, transcription factors (TFs), etc. The activity of any given component (*u*_1_ in this case) makes a pattern. As we show in the text, the dynamics of the activity of any given component can be thought of as motion on a landscape. Minima of the landscape correspond to the entire patterned state. The dynamics on the landscape has a form which is often simple and independent of the details of the underlying regulatory network. A time-independent map from the landscape to the expression of *u*_1_ allows us to quantitatively capture the dynamics of *u*_1_. (B) Biological perturbations like mutants (e.g., deleting a node) or changes to signaling can change the trajectory and lead to a different patterned state. The perturbations can be parameterized in the landscape, resulting in a tilted landscape that takes the same initial point to a pattern with two bumps. The streamplots show changes in the basins of attraction for the pattern with one and two bumps.

**FIG. 2.**
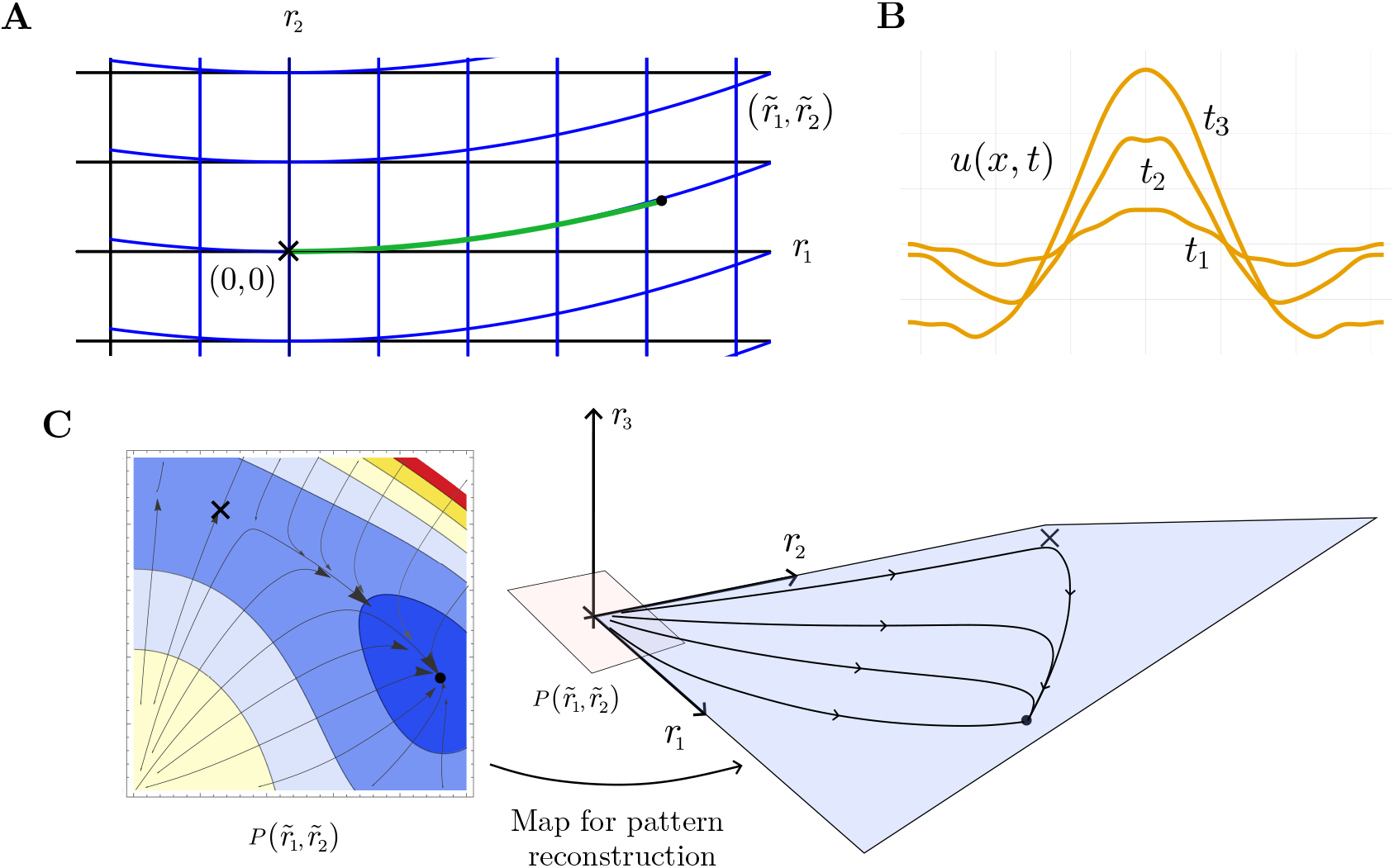
Parameterization of Turing patterns in normal form coordinates with corrections from stable modes. (A) Fourier modes serve as a natural basis for the dynamics of components in models of Turing patterns (rectangular grid shown in black). Here we show plot of the radial part of the dynamics of one unstable (*z*_1_) and one stable (*z*_2_) mode. The initial state (*r*_1_ = *r*_2_ = 0, denoted by cross) is unstable, and the dynamics (trajectory in green) goes to a patterned state (denoted by the filled circle) which generically has a contribution from both modes. We can change coordinates (blue) so that the normal form coordinates align with the dynamics. (B) Spatio-temporal data for the component of interest. (C) We project the data onto the normal form coordinates and fit to the normal form dynamics, which can be expressed as a flow along an effective landscape (shown using contour plot). As the dynamics of the original Fourier modes can be written entirely as a time-independent function of the normal form coordinate, we can reconstruct the dynamics of the component of interest (see text for details).

In general, the coordinate changes involve all of the modes. However, we assume that the dominant contribution to the coordinate changes comes from the unstable modes. By considering all terms allowed by symmetry in the equations for the stable modes and keeping only the ones that involve the unstable modes, we obtain a map for the stable modes. For two modes, one unstable and one stable, the coordinate transformation involves all the non-resonant terms in general

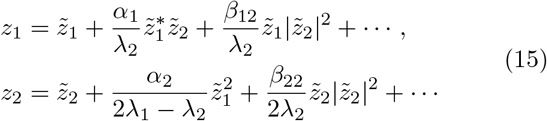

We keep only the dominant terms in the unstable mode 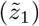, which gives 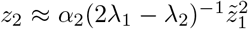. In general, we have to consider the coordinate transformation for all the stable modes we decide to keep. For one unstable mode, the map to stable modes is given by 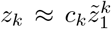, where the parameter *c*_*k*_ is to be fit from the data. Importantly, the coordinate transformation is time-independent. The dynamics of the concentration of any molecule *u*(*x, t*) is thus given by

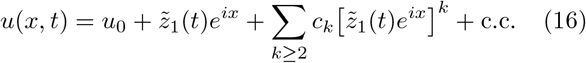

Note that we have absorbed an arbitrary scale in the normal-form variables. The above equation along with Eq. (13) gives us our approximation for the dynamics with only one unstable mode. It is easily generalized to multiple unstable modes.

To test our results, we use a three-component Turing network (Fig. 3A, equations are given in SI Section II A). The network is modeled after the Bmp-Sox9-Wnt network in Ref. [29], but we use Hill functions instead of cubic nonlinearities to model interactions between components.

**FIG. 3.**
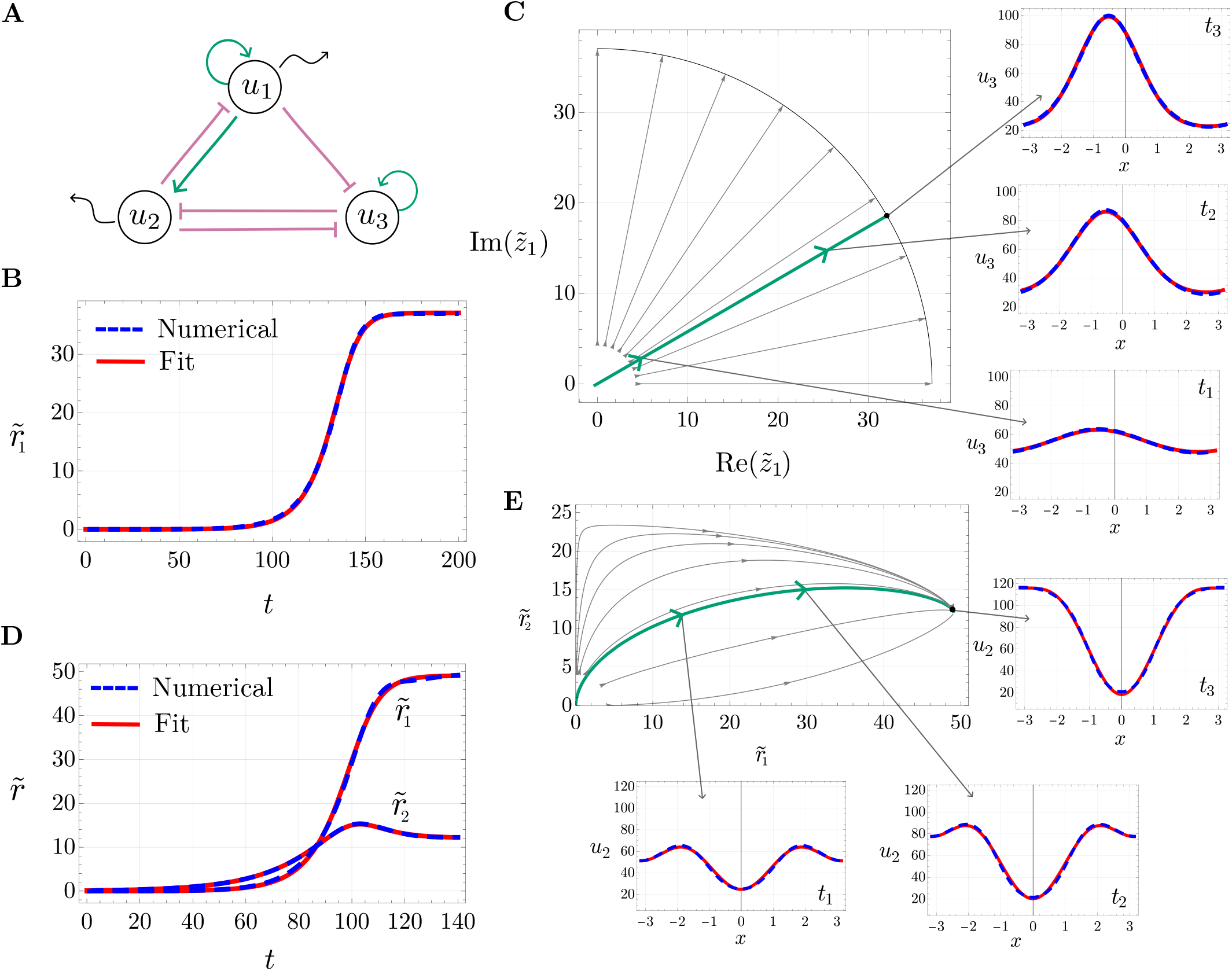
Normal form captures the dynamics of three-component Turing networks. (A) A 3-component network with a non-diffusible component *u*_3_, with activatory (green) and inhibitory (purple) links. (B) Projection of the numerical solution of *u*_3_(*x, t*) obtained by solving the reaction-diffusion equations on the unstable mode (blue, dashed) and fit (red) to the radial part of 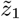 dynamics. (C) The flow of 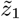 dynamics. Small perturbations away from the origin (homogeneous state) grow radially and stabilize on a ring of fixed points. The angle determines the position of the bump and remains constant. Only one quadrant is shown. The normal form approximation (red) is shown alongside the numerical solution of *u*_3_ (blue, dashed). (D) For two unstable modes, the projection of *u*_2_ dynamics along the unstable modes (blue, dashed) and normal form approximation (red) of the radial part of the dynamics of 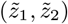. (E) We show the radial flow of the two unstable modes going to a final patterned state. The numerical solution of *u*_2_(*x, t*) (blue, dashed) along with the normal form approximation (red).

For the case of a single unstable mode (*k* = 1), we project the dynamics of the component *u*_3_(*x, t*) along the unstable mode. The angular part of the normal form equation remains constant and the radial part is given simply by 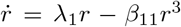. We fit parameters *λ*_1_ and *β*_11_ to the curve obtained from projecting the numerical solution along the unstable mode (Fig. 3B). The parameters of the fit have a simple interpretation. The linear growth rate around the instability is given by *λ*_1_ and the pattern saturates at *r*_0_ = (*λ*_1_*/β*_11_)^1*/*2^. The dynamics of the normal form coordinate (Fig. 3C) goes along the unstable direction towards a circle of fixed points. The angle corresponds to the positioning of the center of the single bump. Fitted parameter values are given in SI Section II A. We show in SI Fig. S2 that including the coordinate changes considered in Eq. (16) is essential to capture the pattern.

For the case of two unstable modes we need to keep both 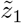 and 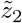 and use the landscape given in Eq. (14b) to approximate the dynamics. We take *θ*_1_ = *θ*_2_ = 0 and fit the numerical solution along the unstable directions to the radial part of the equations derived from the potential given in Eq. (14b). We did not require a metric to fit the dynamics. The fit of the dynamics projected on to the two unstable modes is shown in Fig. 3D. The dynamics of the radial coordinates of the two unstable modes is shown in Fig. 3E. The framework described above is therefore able to capture fairly complex spatio-temporal dynamics.

The normal form of the dynamics of unstable modes can be seen as motion on a potential landscape. Attractors on the landscape correspond to stable patterned states. Biological perturbations to the underlying network effectively modify the landscape parameters, tilting it so the dynamics goes to a different attractor and lead to a different observed pattern (Fig. 1). The observable expression can be systematically recovered from the landscape coordinates. Hence, the landscape is a useful and universal parameterization of the dynamics independent of the details of the underlying networks of interactions.

## IV. PATTERNS ON 2D DOMAIN

We demonstrate how our method can be extended to describe pattern formation on a 2D domain. In 2D, the plane wave *e*^*i****k*.*r***^ is an eigenfunction of the Laplacian ∇^2^ with eigenvalue 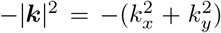 and positional co-ordinate ***r***. Consequently, the growth rate depends only on the magnitude of the wavevector, which allows many combinations of *k*_*x*_ and *k*_*y*_ to become unstable. Nevertheless, these will be discrete on a bounded domain. This allows us to use our formalism for 2D patterns, which we demonstrate on a square domain.

We consider the same reaction-diffusion equations as before, and keep the diffusion constants in the two directions identical (parameter values are given in SI Section II B). With the solution *u*(*x, t*) as the superposition of all the modes, we derive the normal form equations as before. For two unstable modes ***k*** = (1, 0), (0, 1), the normal form is given by

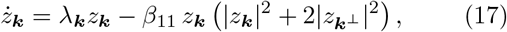

where ***k***^*⊥*^ is the other mode and 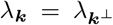. For the stripe pattern shown in Fig. 4A, the instability is only along the *x* direction. Therefore, we ignore the mode corresponding to the *y* direction and obtain the same normal form given in Eq. (13). To simplify things further, we only examine the dynamics of the central stripe by restricting our attention to its neighborhood. The fit to the numerical solution is shown in Fig. 4B. Further, the spatial pattern is now distinctly composed of several modes, but including corrections from a small number of stable modes suffices to give a very good approximation to the full spatio-temporal dynamics (Fig. 4C). Fitted parameter values are given in SI Section II B.

**FIG. 4.**
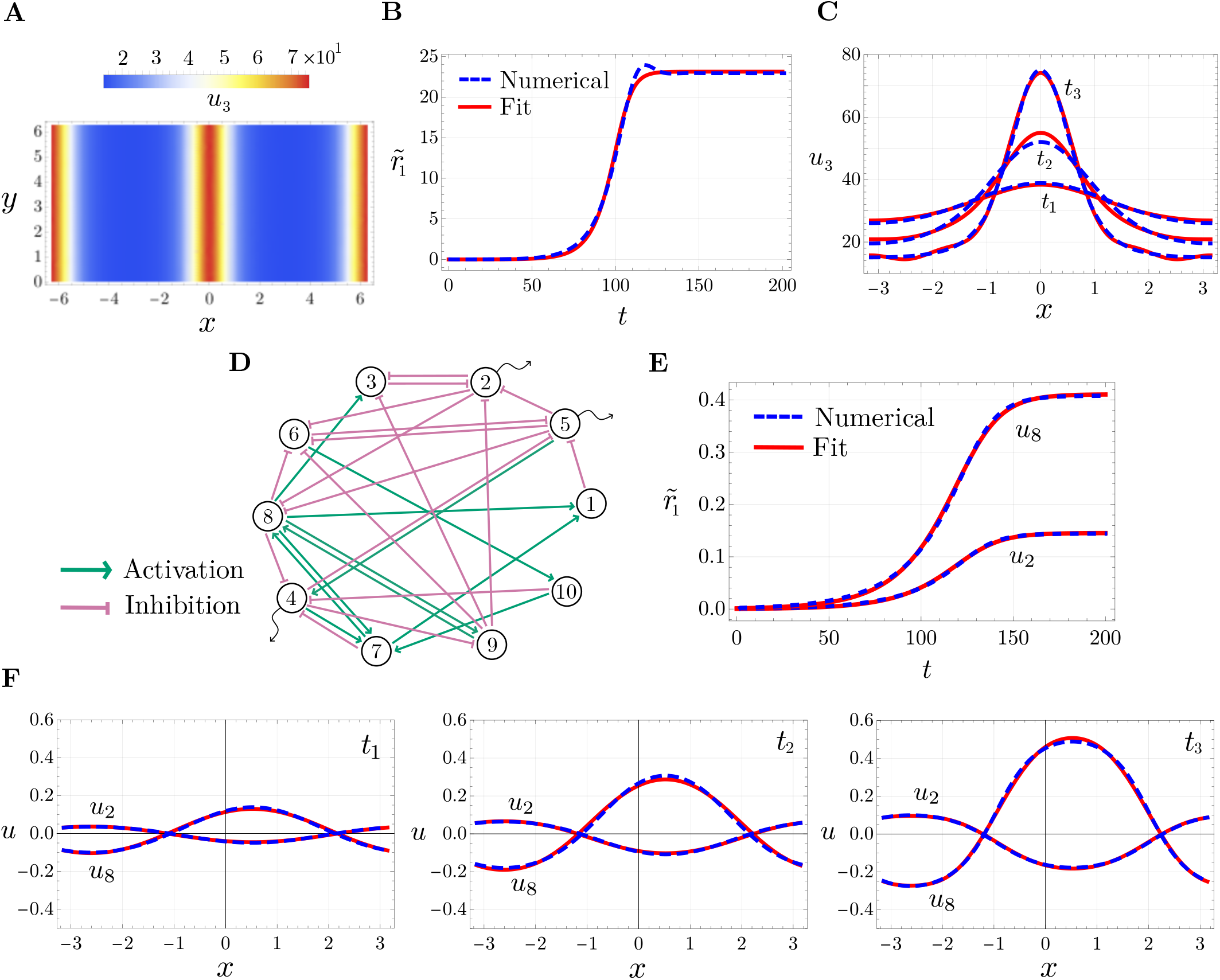
Generalization to patterns on 2D spatial domain and large networks. (A) Numerical solution of the dynamics of *u*_3_(*x, y, t*) on a 2D spatial domain. (B) Projection of the numerical solution of the central stripe along the unstable mode (dotted blue) and the fit (red) to the radial part of the dynamics of 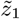. (C) Spatio-temporal dynamics of the central stripe (blue, dashed) with fit (red) to normal form dynamics with corrections from 3 stable modes. (D) A random network with 10 components that are activating or inhibiting. Diffusible components are shown with wiggly arrows. (E) The dynamics of the radial part of the projection of the numerical solution of *u*_2_ and *u*_8_ components along the unstable mode (blue, dashed), along with the normal form approximation (red). (F) Spatio-temporal dynamics of a diffusible component *u*_2_ and non-diffusible component *u*_8_ (blue, dashed) and fit (red) to normal form dynamics with corrections from one stable mode.

## V. PATTERNING DYNAMICS FOR LARGE NETWORKS

The real strength of our formulation is revealed in considering more complex networks. Contrary to the simple networks discussed previously, regulatory and signaling networks are generally complex and involve many interacting components. Even with the addition of more components in the network, the essential structure of the dynamics remains unchanged and is largely independent of the specific details of the individual components. To demonstrate this, we analyze a network with ten components and approximate it with the framework described above.

There is an extensive literature on modeling interactions in complex networks with random matrices [30, 31]. We construct our network (Fig. 4D) by assuming the Jacobian *J* = *G*− *η*𝕀, calculated at the homogeneous state, is a random matrix whose eigenvalues are all negative [32, 33]. The stability is ensured by requiring 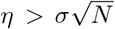, where off-diagonal elements of *G* are taken randomly from a Gaussian distribution and diagonal terms are set to zero (SI Section II C). To saturate the dynamics, we add cubic terms to the equations. With few diffusible components, we search for diffusion constants to give one unstable mode (SI Section II C). Dynamics of a diffusible (*u*_2_) and a non-diffusible (*u*_8_) component along the unstable mode fit to the normal form equation Eq. (13) is shown in Fig. 4E. Despite the complexity of the network, the approximation given by Eq. (13) fits the dynamics well. With corrections from one stable mode, the spatio-temporal dynamics of both the components is shown in Fig. 4F.

## VI. COUPLING POSITIONAL INFORMATION WITH TURING PATTERNS

Recent proposals suggest that Turing patterns can be coupled with positional information to give spatial patterns that may be more robust and reproducible [11, 34]. The resulting dynamics from the coupling of the two mechanisms can be classified depending on whether the spatial eigenfunctions are perturbed. The general form of the equations now becomes

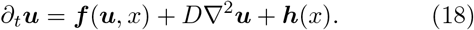

We model a three-component network by directly specifying the Jacobian and use cubic saturation terms (SI Eq. (S7)).

### A. Pattern alignment with production gradient

To study the effect of a production gradient ***h***(*x*), in Eq. (18), we make the reaction terms homogeneous [35]. The production term breaks translational symmetry (Fig. 5A, shown in yellow) but does not change the eigen-functions around the fixed point. For one unstable mode, the normal form equation in radial and angular directions is given by

**FIG. 5.**
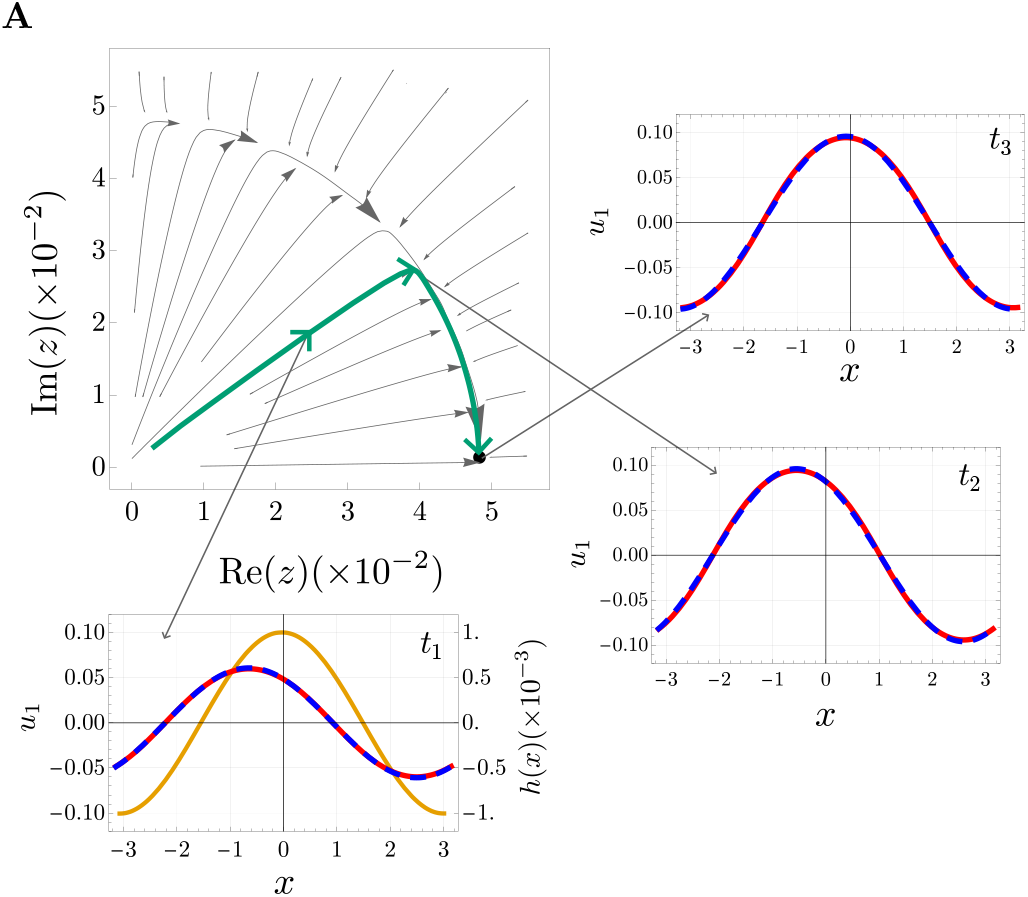
Alignment of Turing patterns with a morphogen gradient. (A) Phase portrait in the 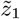 plane for a Turing pattern coupled with a small production gradient, which breaks the translational symmetry and leaves only one fixed point determined by the profile of the morphogen gradient. Spatio-temporal dynamics (blue, dashed) and fit to normal form (red) at time *t*_1_ = 5, *t*_2_ = 20, *t*_3_ = 200. The pattern eventually aligns with the morphogen gradient. The production gradient *h*(*x*) is shown in yellow.

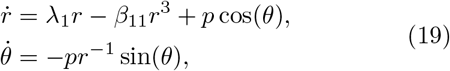

where *p* is the projection of the production gradient along the unstable mode. For *p*≪1, the evolution along the radial and angular directions is separated (Fig. 5A). The pattern grows with a constant phase until saturation, followed by translation, which aligns the bump to a specific position (Fig. 5A). Thus, the production gradient can fix the position of the bump.

Eq. (19) requires a metric to be written in potential form. However, in the limit that we consider, the equations are in potential form as the motion along the two directions is separated. We fit the radial dynamics as done previously in Section III. The solution to the angular dynamics is given by 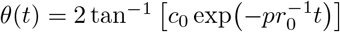, where we have utilized the timescale separation to set *r* to its steady state *r*_0_. We require one parameter to fit the radial dynamics (SI Fig. S3). The spatio-temporal pattern with the fit is shown in Fig. 5A. Parameter values obtained for the fit are given in SI Section II D.

### B. Sequential patterns

A more complex case is given by a wavefront of morphogen traveling at some velocity *v*, which modulates the reaction term ***f*** in Eq. (18). We modulate a parameter with a step function *s*(*x, t*) to give two values, resulting in either the stability of the Jacobian or one unstable mode (SI Section II E). This divides the region into non-patterning and patterning (Fig. 6A), giving us a sequential appearance of bumps behind the moving front (SI Fig. S4). We approximate the wavefront by changing it in discrete steps.

**FIG. 6.**
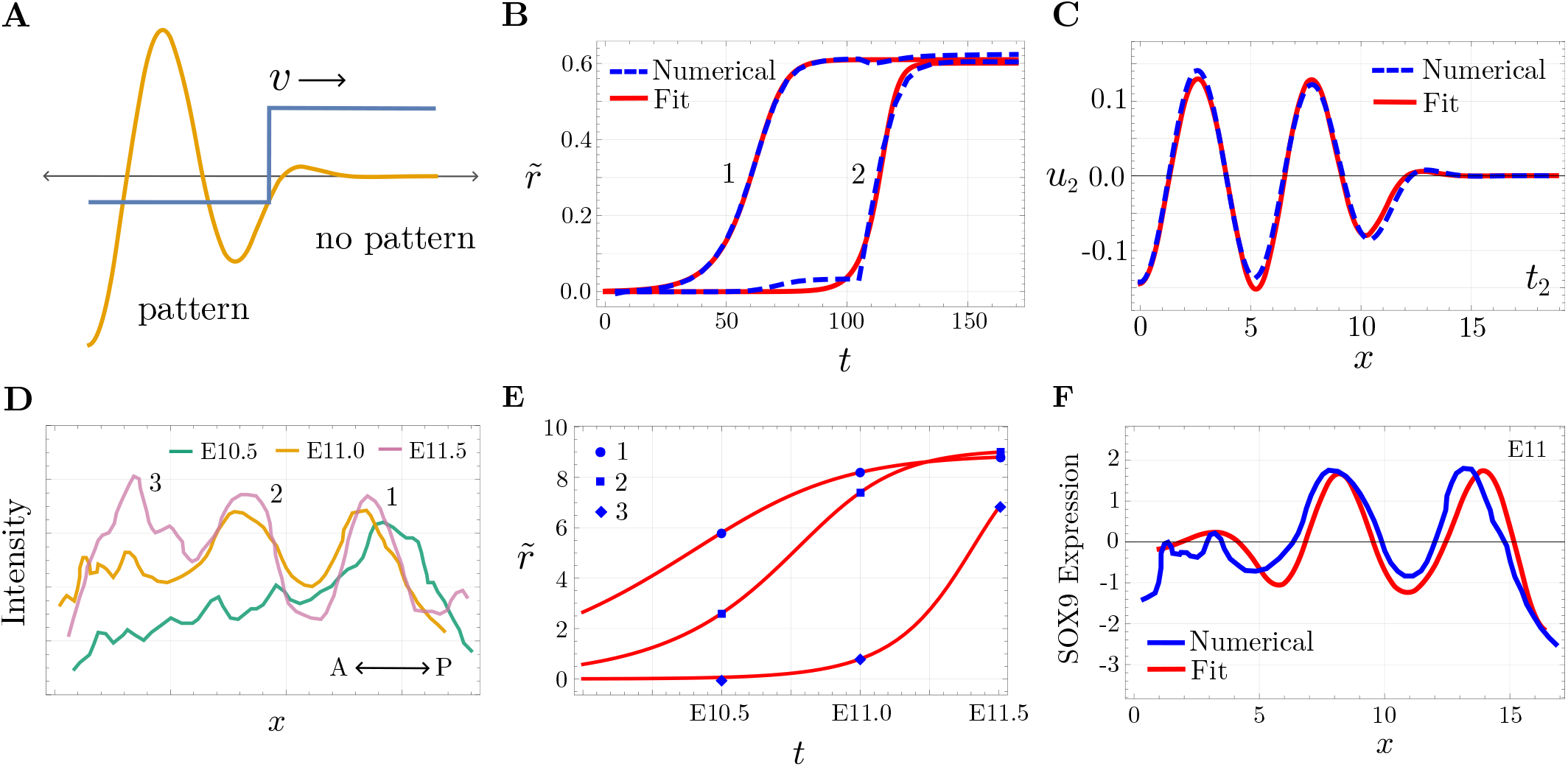
Guided organization of Turing patterns with a modulated step. (A) Step modulation in a parameter of the reaction terms takes the dynamics from a non-patterning regime to a patterning regime. (B) The ‘independent bump’ approximation (see text) fit to the numerical solution. (C) Parameter modulation gives sequential development of pattern (blue, dashed) and fit with the sum of localized eigenfunctions (red) at *t* = 120. (D) Expression of SOX9 along the AP axis at different embryonic time points (data from Fig. 2B in [29]). (E) The ‘independent bump’ approximation fit (red) with experimental data (blue, points). (F) The fit with the sum of localized eigenfunctions (blue) and the SOX9 expression (blue) at E11.

Essentially, each bump grows independently except at the boundary, where continuity requires that the dynamics be connected. We therefore obtain an ‘independent bump’ approximation, and the pattern can be written as a sum of localized eigenfunctions Σ_*i*_ *r*_*i*_(*t*)*ψ*_*i*_(*x*). The dynamics of the amplitudes *r*_*i*_ is fit to the radial part of the normal form equation as before, except the initial condition is an additional fit parameter (Fig. 6B). The profile obtained by the sum of eigenfunctions fits the sequential dynamics well (Fig. 6C, SI Fig. S4). When patterning is slow relative to the front speed, the curves in Fig. 6B move closer and eventually collapse onto each other, recovering Turing patterning with a single effective mode (SI Fig. S4).

Interestingly, during vertebrate limb development, the expression of SOX9 shows such a sequential dynamics [29] (Fig. 6D). We fit the SOX9 data at three time points using the method described above. The fit for each bump is shown in Fig. 6E. Parameter values obtained for the fit are given in SI Section II E. The profile of our fit to the data at time E11.0 (Fig. 6F) is suggestive that this simple approximation may be able to quantitatively capture the data. We note that pattern formation on an apically growing domain also gives similar sequential dynamics and is mathematically similar to modulating a parameter via a front [36, 37].

## VII. DISCUSSION

Turing patterns have received extensive theoretical attention over the years as a model for developmental pattern formation. However, their applicability has been somewhat limited due to the lack of suitable molecular candidates which can implement the proposed mechanisms. Here, we suggest that the observed behaviour of Turing patterns can be directly fit to data. We explicitly calculate the simplest normal form for Turing patterns and show that it can be written as a landscape in normal form coordinates. We also propose a method to reconstruct the observed expression using coordinate changes.

The coordinates of the landscape correspond to Fourier modes and can be derived in a very general way, as we demonstrate. The stable fixed points correspond not to cell fates, but to a particular pattern over many cells. Different fixed points correspond to different possible patterns in the system.

The applicability of our framework relies on the presence of a low-dimensional unstable manifold. It is bound to be useful only if a small number of modes are unstable. There are other instances of patterning (like Notch-Delta patterns), where many unstable directions exist, and the Fourier modes are not the right variables to simplify the dynamics of the system.

Nevertheless, since our framework does not rely on any precise form of the equations, we expect it to be appli-cable to a very large class of systems. For example, the extension to different geometries would be done in an obvious way by using the eigenfunctions of the Laplacian on the given geometry. Similarly, mechanical systems have been shown to have Turing-like instabilities, but the nature of the interactions is not relevant for our framework [38, 39].

We expect our framework to be useful in cases where there are putative Turing patterns alongside time and space-resolved expression of a marker. Knowing the details of all of the regulatory or signaling components is not necessary. Since the behavior we are trying to model is determined by the unstable eigenvector, the dynamics of every component will follow the dynamics of this eigenvector. This leads to a drastic reduction in the number of parameters required to capture the dynamics. For 3-component networks with one unstable mode, the original equations have 18 parameters, whereas the approximation to the dynamics uses only 4 parameters. Moreover, the number of parameters in reaction-diffusion equations increases with the number of components in the equations, so the 10-component network we study technically has 51 parameters, but we need only 3 parameters to approximate the dynamics of any given component. In this sense, our framework constitutes a minimally parameterized description of Turing patterns.

We also demonstrated how the landscape view of Turing patterns changes in the presence of an external morphogen. Earlier studies had pointed out the difference between field and rate modulation of Turing patterns by external morphogens [35]. Mathematically speaking, the modulation by an external component makes the parameters of the reaction-diffusion equation dependent on space. Some kinds of modulations will leave the base spatial eigenfunctions unperturbed and only change their dynamics, as we observe when we introduce a ‘field’ like variable. Others may change the eigenfunctions themselves, as we observe when we change a ‘rate’ like variable.

We discover that the sequential appearance of SOX9 expression in the context of digit formation can be modeled as a Turing pattern coupled with a wavefront of a morphogen. While analytical results are more difficult to obtain in this case, we find that an ‘independent bump’ approximation corresponding to spatially localized eigenfunctions allows us to capture the dynamics. The interaction between these modes may lead to much more complex dynamics. Whether such dynamics can be systematically studied will be a topic for future study. Nevertheless, such a systematic investigation will be of value because it will not rely on a particular form of the equations.

One candidate for such a morphogen in the case of digit formation in mice is Shh [40–42]. Shh perturbations lead to successive digital loss, which would be consistent with our expectations [43, 44], although the precise role of Shh remains under debate [45–47]. If such a modulating morphogen is quantitatively changed, this would be a powerful use of our framework and its predictions.

There are a few different kinds of perturbations that can be parameterized in our framework. One can shift the center of the dispersion relation and change the most unstable mode. This was suggested to be the role of Hox mutants in digit patterning, although the interpretation of these mutants is disputed [48, 49]. More subtle perturbations may only change the timing of pattern formation, which would be modeled by changes in the parameters that describe the normal form flow [50]. The structure of how these parameters depend on possible perturbations needs to be investigated, but previous studies in the case of cell fate have found that simple linear dependence often captures the dynamics quantitatively [22].

Our framework may also be a natural way to investigate the robustness of Turing patterns. The instability conditions are determined at linear order but the higher order terms we consider determine the domain of attraction. The robustness of a pattern depends on the domain of attraction of its attractor in the landscape. Some studies have suggested that the sequential appearance of patterns may be a more robust way to get a fixed number of patterns [34]. It would be interesting to investigate whether this can be shown using our framework.

Our work contributes to the growing theoretical literature on the nonlinear analysis of Turing patterns [51, 52].

However, as mentioned in the Introduction, we distinguish our work from the substantial work on weakly non-linear analysis of Turing patterns using amplitude equations [15, 53]. Our work performs near identity coordinate changes to simplify the equations explicitly and uses the framework of hypernormal form theory. It shows that the dynamics in normal form coordinates can be written as a landscape and makes a specific proposal for how to quantitatively reconstruct the observed expression from these coordinates. Though the reduction to a landscape technically requires a metric, we did not need the metric to fit the data in the cases we considered. We also note that fifth-order terms can not be removed by such coordinate trasformations but we did not need to include them in the cases we investigated. It is possible that such terms and the metric may be needed in case of more complex dynamics [54] or subcritical bifurcations. The experimental data we considered is in the form of static snapshots but time-lapse microscopy would be necessary to distinguish any contribution from the metric.

Ultimately, we believe that our description of Turing patterns will be useful for the study and description of their dynamics in the context of developmental patterns as more quantitative data on gene expression becomes available.

## ACKNOWLEDGEMENTS

We acknowledge support from the Department of Atomic Energy, Government of India (under project RTI4006) and the Simons Foundation (287975). We thank Philip Maini and Tosif Ahamed for their critical reading of a draft of this manuscript and helpful suggestions. SS would like to thank Ankit Dhanuka for helpful discussions.

## Appendix A: Linear stability analysis

Following Ref [55], the characteristic equation det (*J* − *Dk*^2^ − *λ*𝕀) = 0 is written as a cubic equation 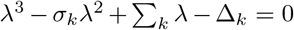 and the coefficients are given by

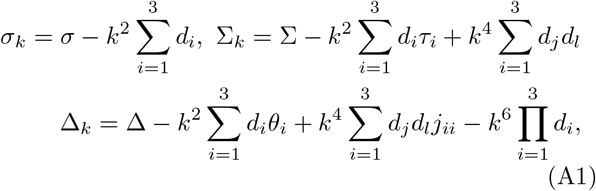

where *σ* = Tr(*J*), Δ = det(*J*), 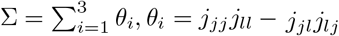 is one of the first minors of *J*, and *τ* = *j*_*jj*_ + *j*_*ll*_ is the trace of the submatrix *J*_*jl*_. The coefficients of the characteristic equation and its roots, are related as follows,

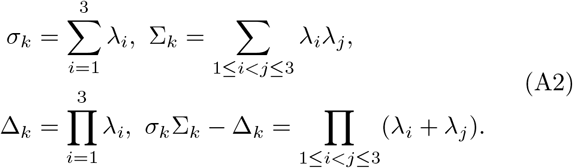

The Routh-Hurwitz criteria gives the following conditions for the homogeneous state to be stable

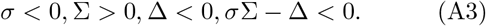

From the above conditions, we have *σ < σ*_*k*_, since *k*^2^ *>* 0 and all diffusion constants are positive. Consequently, in a 2-component system, instability can occur only along one eigenvalue *λ*^(1)^(*k*) since *σ*_*k*_ = *λ*_1_ + *λ*_2_ *<* 0.

We want the homogeneous mode (*k* = 0) and higher modes (*k*→ ∞) to be stable and thus, the eigenvalue *λ*(*k*) changes sign twice. For a 3-component network, having two unstable eigenvalues *λ*^(1)^(*k*) and *λ*^(2)^(*k*), would require four solutions for Δ_*k*_ = 0. However, Δ_*k*_ is a cubic polynomial in *k*^2^, and therefore admits at most three solutions. As a result, only one eigenvalue *λ*^(1)^(*k*) can have an instability.

