## Supplementary material for "A landscape description of the dynamics of Turing patterns": NA

### A landscape description of the dynamics of Turing patterns: Supplementary Information

Shubham Shinde and Archishman Raju
*Simons Centre for the Study of Living Machines*
*National Centre for Biological Sciences, Tata Institute of Fundamental Research, Bangalore, India*
(Dated: April 17, 2026)

#### I. DERIVATION OF NORMAL FORM EQUATIONS FROM REACTION-DIFFUSION EQUATIONS

We consider the two-component activator-inhibitor equations which are given by

$$\begin{aligned}\partial_t a &= \rho + \frac{a^2}{h} - \mu a + d_a \partial_x^2 a \\ \partial_t h &= a^2 - h + d_h \partial_x^2 h\end{aligned} \tag{S1}$$

The homogeneous fixed point is given by $(a_0, h_0) = ((1+\rho)/\mu, (1+\rho)^2/\mu^2)$. For the parameter values

$$\rho = 0.015, \mu = 0.24, d_a = 0.03, d_h = 5$$

modes $k = 1, 2$ become unstable. We Taylor expand the reaction terms around the homogeneous state and keep the nonlinear terms up to cubic order. As discussed in the Main text, to derive the equations for $z_1$ and $z_2$, we substitute the ansatz $(a(x,t), h(x,t))^T = \sum_k \boldsymbol{v}_k z_k e^{ikx}$ in Eq. (S1) and project the resulting expression onto $k$-th mode using $(2\pi)^{-1} \oint \mathrm{d}x e^{ikx}$ followed by dot product with the left eigenvectors corresponding to $\boldsymbol{v}_k$. The equations along $z_1$ and $z_2$ are given by

$$\begin{aligned}\dot{z}_1 &= \lambda_1 z_1 - \alpha_1 z_1^* z_2 - \beta_{11} z_1 |z_1|^2 - \beta_{12} z_1 |z_2|^2, \\ \dot{z}_2 &= \lambda_2 z_2 - \alpha_2 z_1^2 - \beta_{21} z_2 |z_1|^2 - \beta_{22} z_2 |z_2|^2.\end{aligned} \tag{S2}$$

For the parameter values given above, we obtain the following coefficient values for Eq. (S2)

$$\begin{aligned}\lambda_1 = 0.1257, \lambda_2 = 0.0904, \alpha_1 = -0.0505, \alpha_2 = -0.0227, \\ \beta_{11} = 0.0059, \beta_{12} = 0.0102, \beta_{21} = 0.0118, \beta_{22} = 0.0030.\end{aligned}$$

Note there are small differences in $\beta_{12}$ and $\beta_{21}$ and $2\alpha_2$ and $\alpha_1$. These differences come from the weak dependence of the eigenvector $\boldsymbol{v}_k$ on $k$. If $\beta_{12} = \beta_{21}$ and $\alpha_1 = 2\alpha_2$, then these equations can be written as the gradient of a potential.

In this case, we want to write Eq. (S2) as an inverse metric times gradient of a potential. We construct the following potential and the coefficients are derived to give the same fixed points as Eq. (S2)

$$V = \sum_{i=1,2} (-\lambda_i |z_i|^2 + \frac{\beta_i}{2} |z_i|^4) + \alpha \left( z_1^2 z_2^* + z_1^{*2} z_2 \right) + 2\beta |z_1|^2 |z_2|^2. \tag{S3}$$

The origin remains a fixed point with same eigenvalues $(\lambda_1, \lambda_2)$. The saddle point of the flow derived from the potential in Eq. (S3) is given by $z_1 = 0, z_2 = (\lambda_2/\beta_2)^{1/2}$. To match this with the saddle point of Eq. (S2), we obtain $\beta_2 = 0.0031$. Similarly, to match the stable fixed point we obtain parameter values $\alpha = -0.0312, \beta = 0.0075, \beta_1 = 0.00594$.

The metric has to project the flows given by the potential onto the actual dynamics. Following [1], we define the inverse metric as a sum of two operators

$$g^{-1} = \frac{1}{\vec{w}.\vec{z}} (\vec{z}\vec{z}^T + \vec{w}_\perp \vec{w}_\perp^T) \tag{S4}$$

where, $\vec{z} = (\dot{z}_1, \dot{z}_2)$ represents the actual vector flow, $\vec{w}$ is the flow derived from the above potential, and $\vec{w}_\perp$ is the vector perpendicular to $\vec{w}$. The streamplot for the dynamics derived from the potential and the actual dynamics (red) do not match as expected (Fig. S1, left). However, when the potential is multiplied by the above metric, the flow matches the actual dynamics (Fig. S1, right). Technically, this construction breaks at the stable fixed point and requires an independent metric locally, which can be glued to the above construction using a sigmoidal function.

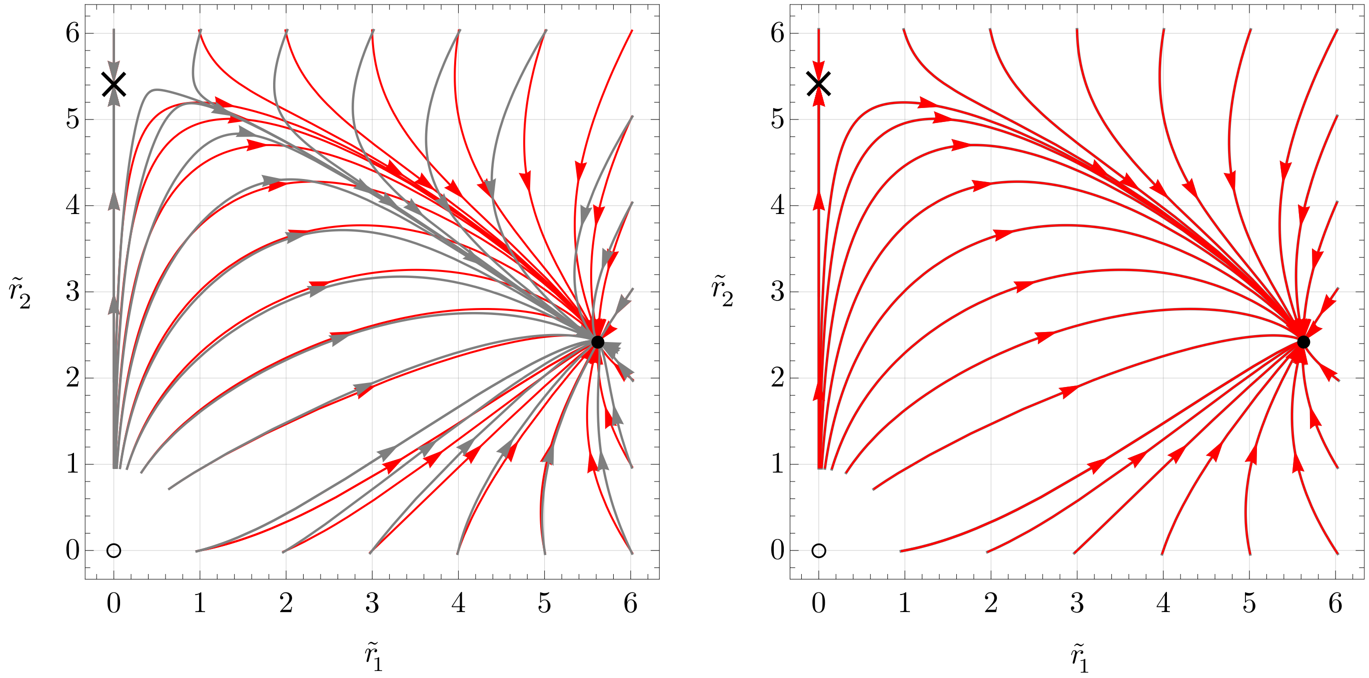


FIG. S1. (Left) Solution trajectories from the actual dynamics (red) and the trajectories of the flow derived from the constructed potential given in Eq. (S3) (gray), (Right) The trajectories of the flow derived by multiplying the potential with the metric (gray) agree with the solution trajectories of the actual dynamics (red).

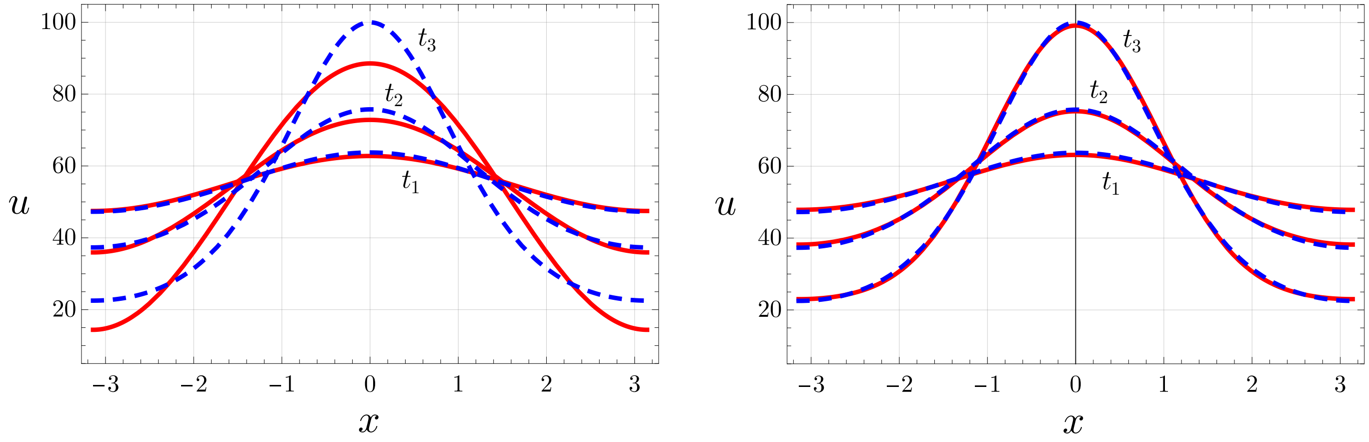


FIG. S2. (Left) Numerical solution at three different times (blue) and reconstruction of the physical solution from only the unstable amplitude (red). The two match at early times but diverge soon. (Right) Same numerical solution (blue) and the reconstruction of the physical solution using the normal form equation and the coordinate change.

However, the deviation of the equation from a potential flow is weak for the reasons mentioned above. Therefore, we did not need the metric in practice to fit the numerical solutions. We also note that fixed points $\boldsymbol{z}_0$ of the system $\dot{\boldsymbol{z}} = g^{-1}(\boldsymbol{z})\nabla V(\boldsymbol{z})$ satisfy $g^{-1}(\boldsymbol{z}_0)\nabla V(\boldsymbol{z}_0) = 0$. Multiplying by $(\nabla V)^T(\boldsymbol{z}_0)$ from the left we get $(\nabla V)^T(\boldsymbol{z}_0)g^{-1}(\boldsymbol{z}_0)\nabla V(\boldsymbol{z}_0) = 0$. Since, $g^{-1}(\boldsymbol{z}_0)$ is positive definite, $(\nabla V)^T(\boldsymbol{z}_0)g^{-1}(\boldsymbol{z}_0)\nabla V(\boldsymbol{z}_0)$ vanishes only when $\nabla V(\boldsymbol{z}_0) = 0$. Therefore, multiplying by a positive definite inverse metric does not alter the fixed points of the system $\dot{\boldsymbol{z}} = \nabla V(\boldsymbol{z})$.

#### II. REACTION-DIFFUSION EQUATIONS AND PARAMETERS

Here, we provide all the parameter values used to get the numerical solutions of reaction-diffusion equations, as well as parameter values obtained for the normal form fit.

##### A. Normal form theory

We model the interactions of the three-component Turing network (Fig. 3A, main text) with Hill functions as

$$\begin{aligned}\partial_t u_1 &= v_1 H_a[k_{11}] H_r[k_{21}] + \nu_1 - \mu_1 u_1 + d_1 \partial_x^2 u_1,\\ \partial_t u_2 &= v_2 H_a[k_{12}] H_r[k_{32}] + \nu_2 - \mu_2 u_2 + d_2 \partial_x^2 u_2,\\ \partial_t u_3 &= v_3 H_r[k_{13}] H_r[k_{23}] H_a[k_{33}] + \nu_3 - \mu_3 u_3,\end{aligned} \tag{S5}$$

where $H_a[k_{ij}] = (1 + (k_{ij}/u_i)^n)^{-1}$ and $H_r[k_{ij}] = (1 + (u_i/k_{ij})^n)^{-1}$ are Hill activation and inhibition functions respectively. Parameter values for the numerical solutions shown in Fig. 3, main text, are given below. For one unstable mode ($z_1$), we use

$$\begin{aligned}&v_1 = v_2 = v_3 = 200, k_{11} = k_{21} = k_{23} = k_{32} = 3, k_{12} = k_{13} = 100, k_{33} = 0.1,\\ &\mu_1 = \mu_2 = \mu_3 = 0.01, \nu_1 = \nu_2 = \nu_3 = 0.1, d_1 = 1, d_2 = 0.01, n = 2\end{aligned}$$

For two unstable modes ($z_1, z_2$), we use

$$\begin{aligned}&v_1 = 70, v_2 = 100, v_3 = 40, k_{11} = 0, k_{21} = k_{23} = k_{32} = 3, k_{12} = k_{13} = 100, k_{33} = 0.0,\\ &\mu_1 = \mu_2 = \mu_3 = 7.6 \times 10^{-3}, \nu_1 = \nu_2 = \nu_3 = 0.09, d_1 = 1, d_2 = 8 \times 10^{-4}, n = 2\end{aligned}$$

We obtain $\lambda_1 = 8.3 \times 10^{-2}, \beta_{11} = 6.06 \times 10^{-5}$ for the normal form fit in Fig. 3B, main text. For the map to the concentration space, we keep two stable modes with coefficients $c_2 = 7 \times 10^{-3}, c_3 = 2 \times 10^{-5}$ (Fig. 3C, main text). The angle $\theta = \pi/6$ is determined from the initial condition. The spatio-temporal pattern in Fig. (3C), main text, is shown at time $t_1 = 120, t_2 = 140$, and $t_3 = 200$.

For two unstable modes, we fit the radial dynamics with $\lambda_1 = 9.2 \times 10^{-2}, \lambda_2 = 5.5 \times 10^{-2}, \beta = 2.9 \times 10^{-5}, \gamma = -5.3 \times 10^{-4}, \beta_1 = 4.0 \times 10^{-5}, \beta_2 = 9.4 \times 10^{-5}$ (Fig. 3D, main text). We keep one stable mode $k_3 = c_1 \tilde{z}_1^3 + c_2 \tilde{z}_1 \tilde{z}_2$ and parameter values $c_1 = -2 \times 10^{-5}, c_2 = 4 \times 10^{-3}$ to fit the pattern in Fig. 3E, main text. The spatio-temporal pattern in Fig. (3E), main text, is at time $t_1 = 90, t_2 = 100$, and $t_3 = 130$.

##### B. Pattern on 2D domain

We use the same reaction-diffusion equations used in the previous section. Parameter values used to obtain numerical solutions in Fig. 4A, main text, are

$$\begin{aligned}&v_1 = 70, v_2 = 100, v_3 = 40, k_{11} = 0, k_{21} = k_{23} = k_{32} = 3, k_{12} = k_{13} = 100, k_{33} = 0.0,\\ &\mu_1 = \mu_2 = \mu_3 = 7.6 \times 10^{-3}, \beta_1 = \beta_2 = \beta_1^{(1)} = 0.09, d_1 = 1, d_2 = 1.5 \times 10^{-3}, n = 2\end{aligned}$$

We obtain $\lambda_1 = 0.105, \beta_{11} = 1.95 \times 10^{-4}$ for the normal form fit (Fig. 4B). We keep three stable modes with coefficient values $c_1 = 0.024, c_2 = 4.9 \times 10^{-4}, c_3 = 9 \times 10^{-6}$ for the map onto concentration space shown in Fig. 4C, main text. The spatio-temporal pattern is fit at time $t_1 = 90, t_2 = 120, t_3 = 150$, Fig. 4C, main text.

##### C. Large network

The reaction-diffusion equation for each component $u_i$ is given by

$$\partial_t u_i = \sum_j J_{ij} u_j + \mu u_i^2 - \nu u_i^3 + d_i \partial_x^2 u_i, \tag{S6}$$

We construct the Jacobian matrix as $J = G - \eta \mathbb{I}$, where $G$ is a random matrix with elements $g_{ij}$ taken from a Gaussian distribution with zero mean and variance $\sigma^2$, and diagonal elements are set to zero. With $\sigma^2 = 0.01, \eta = 2$, all the

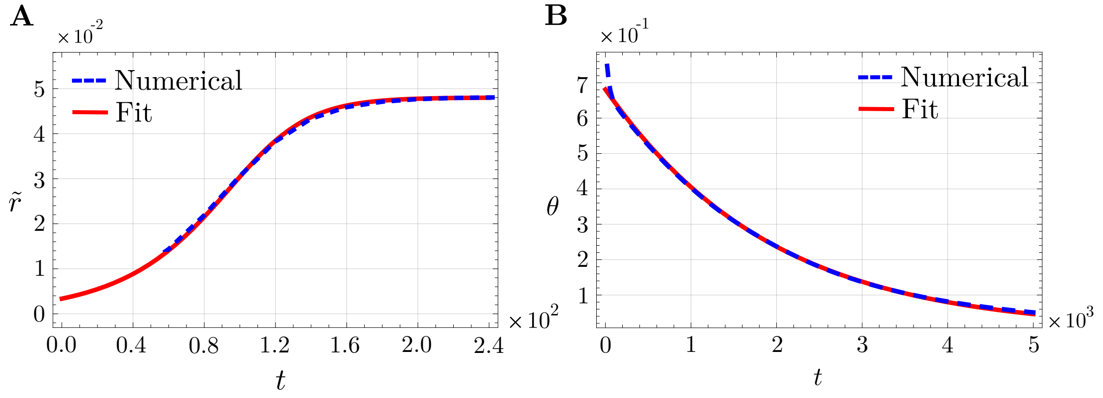


FIG. S3. Dynamics along the radial and angular directions for one unstable mode.

modes are stable. We keep diffusion in three components, and the diffusion constants are sampled from a uniform distribution over an interval of $[0, 20]$ to make one mode $(z_1)$ unstable. We get $d_2 = 0.05, d_4 = 1.0, d_5 = 2.0$ and use identical parameter values $\mu = 0.02, \nu = 0.05$ for each component. The Jacobian matrix used to get the numerical solution in Fig. 4E and F, main text, is

$$J = \begin{bmatrix} -0.60 & 0.00 & 0.00 & 0.00 & -0.01 & 0.00 & 0.00 & 0.00 & 0.00 & 0.00 \\ 0.00 & -0.60 & -0.09 & 0.00 & 0.00 & -0.19 & 0.00 & -0.22 & 0.00 & 0.00 \\ 0.00 & -0.09 & -0.60 & 0.00 & 0.00 & 0.00 & 0.00 & 0.00 & 0.00 & 0.00 \\ 0.00 & 0.00 & 0.00 & -0.60 & -0.32 & 0.00 & 0.13 & 0.00 & -0.12 & 0.00 \\ 0.00 & -0.09 & 0.00 & 0.05 & -0.60 & -0.17 & 0.00 & -0.35 & 0.00 & 0.00 \\ 0.00 & 0.00 & 0.00 & 0.00 & -0.01 & -0.60 & 0.00 & 0.00 & 0.00 & 0.10 \\ 0.15 & 0.00 & 0.00 & -0.13 & 0.00 & 0.00 & -0.60 & 0.21 & 0.00 & 0.00 \\ 0.21 & 0.00 & 0.06 & -0.14 & 0.00 & -0.19 & 0.26 & -0.60 & 0.43 & 0.00 \\ 0.00 & -0.46 & -0.16 & 0.00 & 0.00 & -0.02 & 0.00 & 0.57 & -0.60 & 0.00 \\ 0.00 & 0.00 & 0.00 & -0.03 & 0.00 & 0.00 & 0.10 & 0.00 & 0.00 & -0.60 \end{bmatrix}.$$

Parameters obtained for the normal form fit for both the components $u_2$, and $u_8$ are $\lambda_1 = 0.045, \beta_{11} = 2.13$ and $\lambda_1 = 0.045, \beta_{11} = 0.26$ respectively (Fig. 4E, main text). We keep two stable modes corresponding to $z_2$ and $z_3$ modes. We use $c_2 = -0.4, c_3 = -2.0$ for $u_2$ and $c_2 = 0.15, c_3 = -0.3$ for $u_8$ to fit the pattern in Fig. 4F, main text. The spatio-temporal pattern is fit at time $t_1 = 100, t_2 = 120$, and $t_3 = 200$, Fig. 4F, main text.

##### D. Pattern alignment with production gradient

To study the effect of positional information on reaction-diffusion equations, we use the following equations

$$\begin{aligned} \partial_t u_1 &= k_{11}u_1 + k_{31}u_3 - u_1^3 + d_1\partial_x^2 u_1 + h(x), \\ \partial_t u_2 &= k_{22}u_2 + k_{32}u_3 - u_2^3 + d_2\partial_x^2 u_2, \\ \partial_t u_3 &= k_{13}(1 - s(x,t))u_1 + k_{23}u_2 + k_{33}u_3 - u_3^3. \end{aligned} \tag{S7}$$

we have added a production gradient $h(x)$ in the equation for $u_1$ and a parameter gradient by modulating the parameter $k_{13}$ as $k_{13}(1 - s(x,t))$. The parameter values used are

$$\begin{aligned} &k_{11} = 1, k_{31} = 1, k_{22} = 1, k_{32} = 1, k_{13} = 0.25, \\ &k_{23} = -0.25, k_{33} = -0.05, d_1 = 1, d_2 = 4 \end{aligned}$$

To investigate the effect of the production gradient, we set $s(x,t) = 0$ and take $h(x) = 10^{-3}\cos(x)$. We obtain $\lambda_1 = 0.49, \beta_{11} = 213.14$ to fit the radial dynamics to the numerical solution along the unstable mode. We use $p = 5.23 \times 10^{-4}$ to fit the angular dynamics. The spatio-temporal pattern in Fig. (5A), main text, is fit at time $t_1 = 5, t_2 = 20, t_3 = 200$. The plots for the radial and angular dynamics with the fit are shown in Fig. S3.

##### E. Sequential dynamics

To get a sequential appearance of bumps, in Eq. (S7), we set the production gradient $h(x)$ to zero and take $s(x,t) = x - 2\pi \lceil t/T \rceil$. $v = 2\pi/T$ is the effective velocity of the front. Parameter values used to get the sequential dynamics shown in Fig. S4A (blue, dashed) are

$$k_{11} = 2.8, k_{22} = 0.8, k_{31} = 1, k_{32} = 1, k_{13} = 0.25$$
$$k_{23} = -0.4, k_{33} = -0.01, d_1 = 3, d_2 = 15, v = 5.23 \times 10^{-2}$$

As discussed in the main text, the dynamics is fit to the sum of eigenfunctions $u_2(x,t) = \sum_i r_i(t)\psi_i(x)$. The eigenfunctions are calculated numerically with a time-independent function $s(x,\xi) = \Theta(-x+\xi) - \Theta(x - 2\pi - \xi)$, and $\xi$ controls the position of the function. The shift $\xi_i$ used to calculate the eigenfunctions is a parameter to be fit, and we use $\xi_1 = 0$ and $\xi_2 = 1.63\pi, \xi_3 = 3.3\pi$ to get the localized eigenfunctions (Fig. S5).

At each time point, the numerical solution $u_2(x,t)$ is fit to $\sum_i c_i(t)\psi_i(x)$. The amplitudes $r_i(t)$ are taken to follow the radial dynamics of the normal form equation. The fitting procedure is the same as before, but the initial condition has to be treated as a parameter to fit. We use $\lambda_1 = 0.091, \beta_{11} = 0.24, c_0 = 1.3 \times 10^{-3}$ and $\lambda_1 = 0.16, \beta_{11} = 0.45, c_0 = 2.66 \times 10^{-9}$ to fit dynamics of the amplitude of the first and second bumps respectively (Fig. 5C, main text). As the front velocity is increased, the two amplitude dynamics curves come closer (Fig. S4B) and eventually overlap in the limit of high velocity (Fig. S4C). For the front velocity $v = 2\pi/120$, we obtain $\lambda_1 = 7.6 \times 10^{-2}, \beta_{11} = 2.64, c_0 = 7.33 \times 10^{-4}$ to fit the amplitude of the first bump and $\lambda_1 = 8.57 \times 10^{-2}, \beta_{11} = 3.17, c_0 = 7.46 \times 10^{-5}$ for the second bump (Fig. S4B).

In the limit of high front velocity, all the bumps emerge simultaneously, and hence the dynamics for the amplitudes $r_i$ is the same for all the eigenfunctions. Thus, we require only one set of parameters $\lambda_1 = 8.72 \times 10^{-2}, \beta_{11} = 3.51, c_0 = 4.19 \times 10^{-5}$ and the fit to the pattern is shown in Fig. S4C.

###### *SOX9 dynamics*

We fit the dynamics of SOX9 to $\sum_{i=1}^{3} r(t)\psi_i(x)$ and follow the same procedure as in the previous section. The eigenfunctions are calculated with the same parameters as before, except for the value of the shift $\xi$. We use values $\xi_1 = 0.6, \xi_2 = 1.6\pi$, and $\xi_3 = 3.57\pi$.

The average intensity of SOX9 expression grows asymmetrically in the two halves of the domain. Since our model is built to capture the dynamics of patterns around a homogeneous state, we exclude the left half of the data at E10.5 while fitting and subtract the mean intensity calculated over the right half of the data. At other timepoints, we subtract the mean intensity calculated over the entire domain.

The dynamics of each amplitude, which follows the radial part of the normal form equation, is fit to coefficients obtained by fitting the sum of eigenfunctions to the SOX9 expression data at each time point. The normal form fit for the three bumps is shown in Fig. 5F. We use parameters $\lambda_1 = 1.01, \beta_{11} = 1.27 \times 10^{-2}, c_0 = 1.001$ for the first bump, $\lambda_1 = 1.54, \beta_{11} = 1.86 \times 10^{-2}, c_0 = 0.12$ for the second bump, and $\lambda_1 = 2.56, \beta_{11} = 3.15 \times 10^{-2}, c_0 = 3.57 \times 10^{-4}$ for the third bump. With the above parameter values, the independent bump fit to SOX9 data is shown in Fig. S7.

[1] David A Rand, Archishman Raju, Meritxell Sáez, Francis Corson, and Eric D Siggia. Geometry of gene regulatory dynamics. *Proceedings of the National Academy of Sciences*, 118(38):e2109729118, 2021.

[2] Jelena Raspopovic, Luciano Marcon, Laura Russo, and James Sharpe. Digit patterning is controlled by a bmp-sox9-wnt turing network modulated by morphogen gradients. *Science*, 345(6196):566–570, 2014.

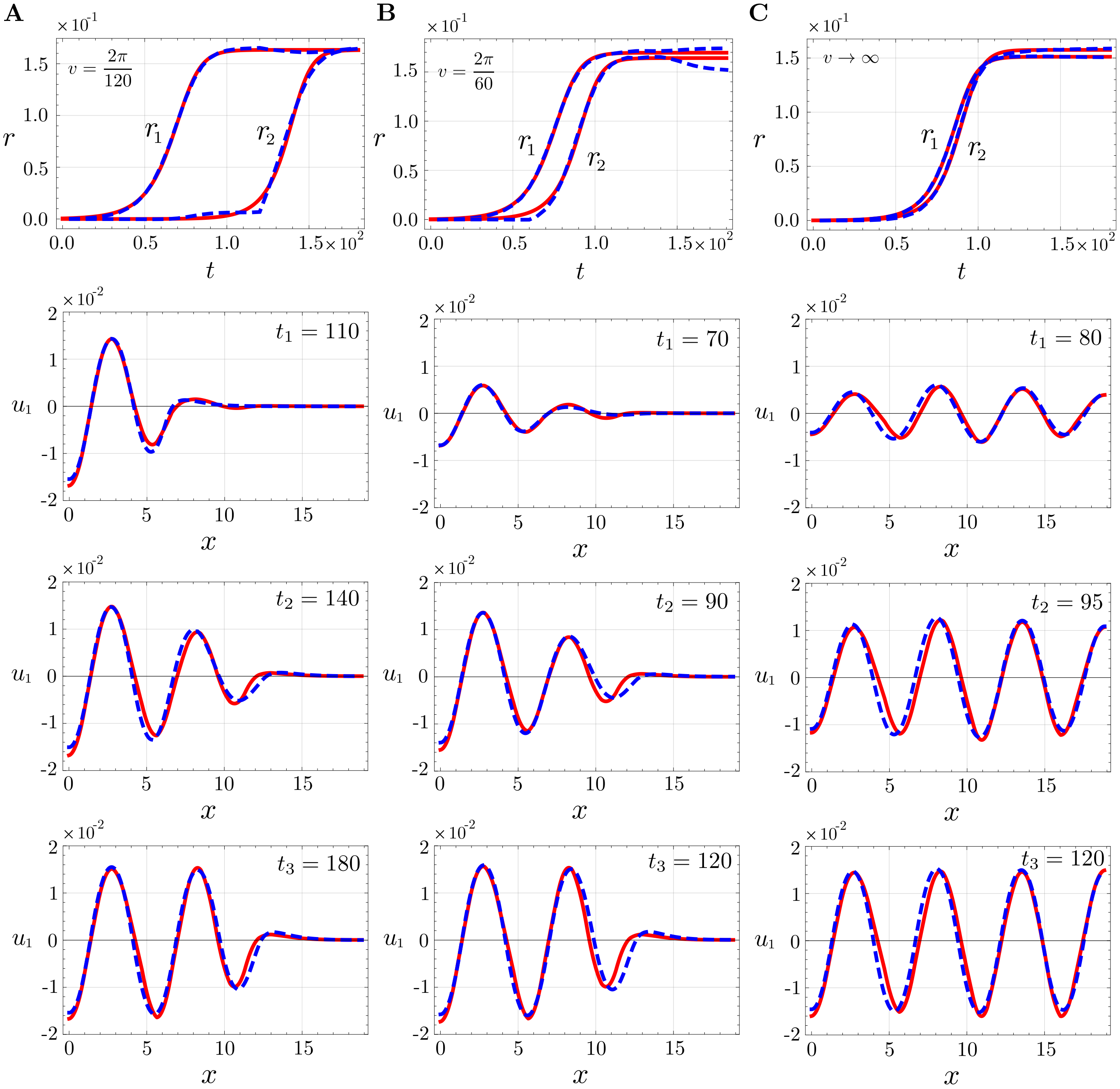


FIG. S4. (A) The dynamics of $r_1$ and $r_2$ from numerical solution (blue, dotted) and fit (red) with the radial part of the normal form dynamics. Pattern at different times (blue, dotted) and fit as the sum of eigenfunctions (red). We focus primarily on capturing the fast dynamics of the pattern formation and ignore the slower dynamics associated with minor rearrangements of the pattern. (B, C) Same as above for $v = 2\pi/60$ and $v \to \infty$.

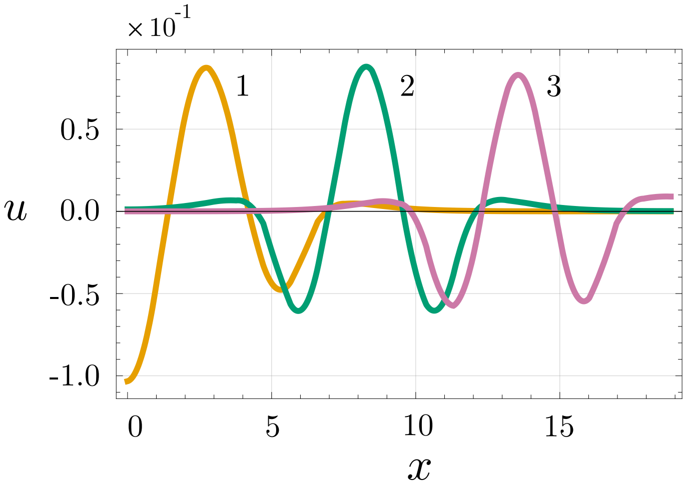


FIG. S5. Eigenfunctions with localized box function for different shift values of $\xi$.

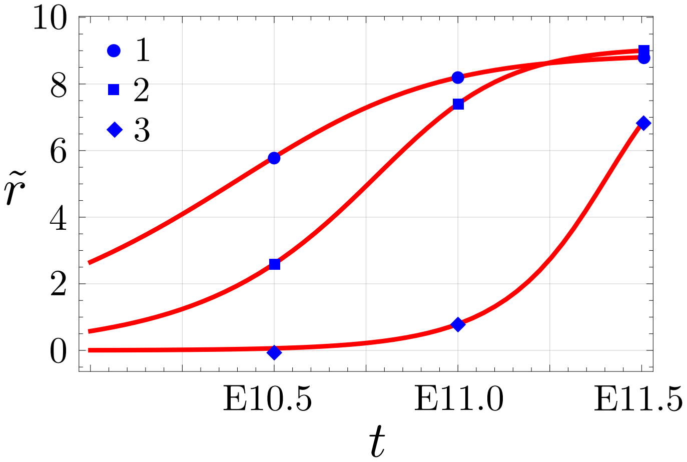


FIG. S6. The normal form fit to the dynamics of amplitudes corresponding to each bump in SOX9 expression, data (blue, points) and fit (red).

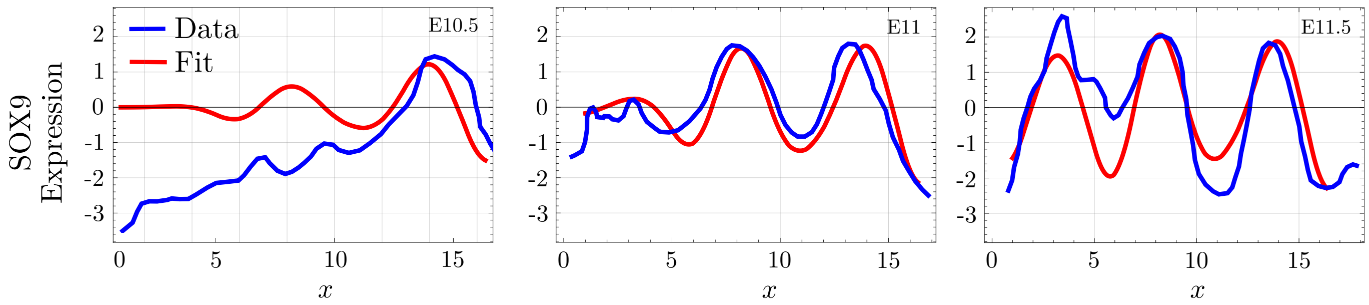


FIG. S7. The expression of SOX9 data (from Ref. [2]) (blue) and independent bump approximation fit (blue) at three different time points.
